# Root hydraulic properties: an exploration of their variability across scales

**DOI:** 10.1101/2023.12.06.570353

**Authors:** Juan C. Baca Cabrera, Jan Vanderborght, Valentin Couvreur, Dominik Behrend, Thomas Gaiser, Thuy Huu Nguyen, Guillaume Lobet

## Abstract

Root hydraulic properties are key physiological traits that determine the capacity of root systems to take up water, at a specific evaporative demand. They can strongly vary among species, cultivars or even within the same genotype, but a systematic analysis of their variation across plant functional types (PFTs) is still missing. Here, we reviewed published empirical studies on root hydraulic properties at the segment-, individual root-, or root system scale and determined its variability and the main factors contributing to it.

We observed an extremely large range of variation (of orders of magnitude) in root hydraulic properties, but this was not caused by systematic differences among PFTs. Rather, the (combined) effect of factors such as root system age, driving force used for measurement, or stress treatments shaped the results. We found a significant decrease in root hydraulic properties under stress conditions (drought and aquaporin inhibition) and a significant effect of the driving force used for measurement (hydrostatic or osmotic gradients). Furthermore, whole root system conductance increased significantly with root system age across several crop species, causing very large variation in the data (> 2 orders of magnitude). Interestingly, this relationship showed an asymptotic shape, with a steep increase during the first days of growth and a flattening out at later stages of development. This behaviour was also observed in simulations with computational plant models, suggesting common patterns across studies and species.

These findings provide better understanding of the main causes of root hydraulic properties variations observed across empirical studies. They also open the door to better representation of hydraulic processes across multiple plant functional types and at large scales. All data collected in our analysis has been aggregated into an open access database (https://roothydraulic-properties.shinyapps.io/database/), fostering scientific exchange.

## 1 Introduction

Root water uptake is a fundamental mechanism essential for the survival of plants. The ability of plants to absorb water through their roots and transport it to the plant’s above-ground tissues is crucial for enabling key physiological processes such as photosynthesis, nutrient absorption, and cell expansion (Lambers & Oliveira, 2019). The effectiveness of root systems in absorbing water allows plants to regulate their water balance, postpone or avoid water stress, regulate canopy temperature, and sustain physiological functions at their optimum (Steudle, 2000a; Lynch *et al*., 2014; Abdalla *et al*., 2022).

Water uptake is a passive process driven by the water potential gradients in the soil-plant-atmosphere continuum (catenary process, Cowan, 1965), where water is pulled up from the soil into the root xylem and up to the leaf following the cohesion-tension principle (Steudle, 2001). Water flow through the root system can be described analogously to electric current through a network of resistances (Landsberg & Fowkes, 1978). The water flow rate (*J,* m^3^ s^-1^) between any two points is dependent on the water potential difference (𝜓, MPa) and the hydraulic conductance (*K*, m^3^ MPa^-1^ s^-1^, the inverse of a resistance) between these points. In that, root water uptake from the root-soil interface to the above ground organs is affected by root hydraulic properties (the individual resistances) and the root system architecture (the way resistances are connected to form a network) (Doussan *et al*., 1998; Leitner *et al*., 2014; Lobet *et al*., 2014) (Figure 1).

**Figure 1:**
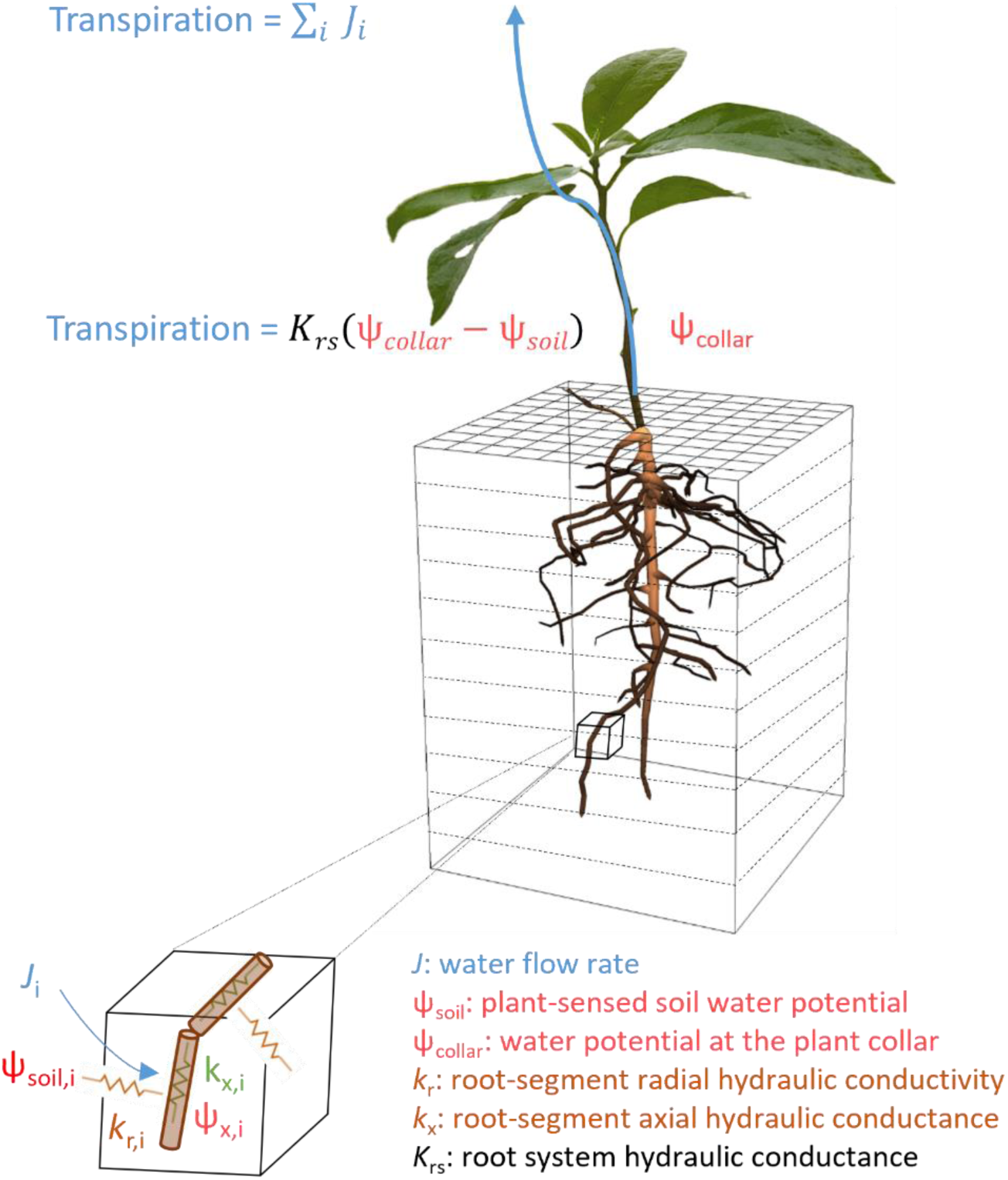
Root hydraulic properties and water flow in the soil-plant-atmosphere continuum. Figure adapted from Vanderborght *et al*. (2021)

Root hydraulic properties can be expressed at different tissue scales, from root segments up to the whole root system (Figure 1, Table 1). The radial conductivity (*k*_r_) represents the capacity of roots to transport water from the root-soil interface to the root-xylem across their radial pathways, and depends on several anatomical features (Steudle, 2000a; North & Peterson, 2005) and aquaporin expression (Gambetta *et al*., 2017). The axial conductance *k*_x_ refers to the ability of roots to transport water longitudinally, which is a function of the number and diameter of xylem vessels (Hacke & Jansen, 2009). The resulting total conductivity of individual roots or root segments (*k*_root_) can be limited by its radial (Bramley *et al*., 2009) or axial components (Sanderson *et al*., 1988; Bouda *et al*., 2018; Boursiac *et al*., 2022a). The whole root system conductance (*K*_rs_) integrates the contribution of all individual conductances along the root system, i.e., it depends on *k*_r_ and *k*_x_ (Bouda *et al*., 2018; Meunier *et al*., 2019) but also on the root system architecture (Doussan *et al*., 2006), and reflects the overall hydraulic efficiency of the root system in transporting water from the soil to the above-ground tissues (see Table 1 for details). Understanding the variability in these key hydraulic properties among and within plant species and in response to changing environmental conditions and environmental stresses is essential for the study of plant water relations (Gallardo *et al*., 1996; Lambers & Oliveira, 2019).

**Table 1:** Root hydraulic properties definitions.

| Symbol | Definition | Tissue level | Units | Alternative symbols used in the literature | Specification |
| --- | --- | --- | --- | --- | --- |
| $k_r$ | Radial hydraulic conductivity | Individual roots or root segments | $\text{m MPa}^{-1} \text{ s}^{-1}$ | $L_r$ (Huang & Nobel, 1994; North & Peterson, 2005; Doussan *et al.*, 2006) | Usually not directly measured, but calculated using $k_{\text{root}}$ and $k_x$ measurements, based on the model of Landsberg & Fowkes (1978). |
| $k_x$ | Specific axial hydraulic conductance | Individual roots or root segments | $\text{m}^4 \text{ MPa}^{-1} \text{ s}^{-1}$ | $K_h$ (Huang & Nobel, 1994; North & Peterson, 2005; Doussan *et al.*, 2006); $K_x$ (Ahmed *et al.*, 2018); $L_x$ (Frensch & Steudle, 1989; Melchior & Steudle, 1993) | The ability of roots to transport water longitudinally |
| $k_{x\_cs}$ | $k_x$ normalized by cross sectional area | Individual roots or root segments | $\text{m}^2 \text{ MPa}^{-1} \text{ s}^{-1}$ | $K_s$ (Pratt *et al.*, 2007; Choat *et al.*, 2012) | $k_x$ data for woody species is very commonly reported on a cross sectional area basis (sapwood, stele, total root cross section) |
| $k_{\text{root}}$ | (Total) root hydraulic conductivity | Individual roots or root segments | $\text{m MPa}^{-1} \text{ s}^{-1}$ | $L_{\text{pr}}$ (Steudle, 2000a; Kim *et al.*, 2018; Boursiac *et al.*, 2022b); $L_p$ (Huang & Nobel, 1994; North & Peterson, 2005; Gambetta *et al.*, 2017; Lambers & Oliveira, 2019) | The total water transport capacity of an individual root or a root segment. It can be separated into its radial and axial components. Often assumed to be an approximation of $k_r$ in the literature (i.e. water transport only limited by $k_r$, not by $k_x$) |
| $K_{rs}$ | Whole root system conductance | Entire root system | $\text{m}^3 \text{ MPa}^{-1} \text{ s}^{-1}$ | $K_{\text{root}}$ (Cai *et al.*, 2022); $L_p$ (Lambers & Oliveira, 2019); $L_{\text{pr}}$ (Steudle, 2000a; Kim *et al.*, 2018), $L_0$ (Maurel *et al.*, 2010; Tyerman *et al.*, 2017; Boursiac *et al.*, 2022b) | The water transport capacity of the entire root system. |
| $K_{rs\_norm}$ | $K_{rs}$ normalized by a measure of the root system size | Entire root system | Depends on normalization | | Most common normalizations found in the literature include:<br>Root surface area: $K_{rs\_area}$ ($\text{m MPa}^{-1} \text{ s}^{-1}$)<br>Root fresh or dry weight: $K_{rs\_weight}$ ($\text{m}^3 \text{ MPa}^{-1} \text{ s}^{-1} \text{ g}^{-1}$)<br>Root length: $K_{rs\_length}$ ($\text{m}^3 \text{ MPa}^{-1} \text{ s}^{-1} \text{ m}^{-1}$)<br>Root volume: $K_{rs\_vol}$ ($\text{m}^3 \text{ MPa}^{-1} \text{ s}^{-1} \text{ m}^{-3}$) |

A large range of empirical methods has been developed for the determination of root hydraulic properties, from the cell and tissue level (Steudle, 1990) up to the whole root system (Tyree *et al*., 1995), with the pressure chamber, the High Pressure Flow Meter (HPFM) and root exudation being the most common ones (Boursiac *et al*., 2022b). While these methods rely on the direct measurement of water flow across root tissues, also more indirect methods based on observations of soil water content and transpiration changes in combination with modelling have been applied (Abdalla & Ahmed, 2021; Abdalla *et al*., 2022). However, different measurement methods may produce different results, especially when comparing methods that rely on a hydrostatic driving force for water flow against those using an osmotic one (Kim *et al*., 2018). Additionally, empirical studies have shown that root hydraulic properties can strongly vary (up to orders of magnitude) among species (Steudle, 2000a; Bramley *et al*., 2009; Pratt *et al*., 2010), but also among genotypes of one species (Rishmawi *et al*., 2023) or even among individuals of the same genotype (Steudle, 2000a). This large variability can be explained, at least partially, by the function of roots as hydraulic rheostats, i.e., the dynamic changes that root hydraulic properties undergo during development and in response to environmental stimuli (Maurel *et al*., 2010). Interestingly, though, a systematic study of the range of variability of root hydraulic properties across multiple plant functional types (PFTs), experimental treatments and measurement techniques is still missing. PFTs provide a simplified description of plant diversity, facilitating the representation of ecosystem processes and vegetation dynamics (Wullschleger *et al*., 2014). Understanding the variability of root hydraulic properties among and within PFTs is therefore key for a better modelling representation of root water uptake processes across scales (Sulis *et al*., 2019; Nguyen *et al*., 2020; Nguyen *et al*., 2022).

In this context, the present study focused on improving the understanding of the variability of root hydraulic properties observed across species and PFTs. For this, we systematically reviewed published empirical studies and addressed the following questions: (i) what is the total range of variation in root hydraulic properties observed in the literature?; (ii) are there systematic differences in root hydraulic properties among PFTs and which other factors affect root hydraulic properties variability?; (iii) are the responses of root hydraulic properties to environmental stresses consistent across PFTs?; and (iv) how are root hydraulic properties affected by root development (root age)?

Given the large amount of data obtained in the review and its complexity (see 2.2 for a detailed data description), the results presented in this study have a stronger focus on *K*_rs_, a key trait that might determine the water use of plants under changing environmental conditions (Vadez, 2014) and integrates the variability of *k*_r_, *k*_x_ and root architecture. But, all original data that was collected in the review has been aggregated to an open access database, which can be easily accessed through a web application (Baca Cabrera, 2023), facilitating data access and further use. Furthermore, we complemented our review by using functional-structural modelling, to improve our understanding of the mechanisms behind the emerging patterns in the empirical data.

## 2 Methods

### 2.1 Literature review selection criteria

The main goal of this study was to obtain an overview about the range of variation in root hydraulic properties observed experimentally, and the main factors contributing to it. For this, we reviewed scientific articles in which whole root system hydraulic conductance, root hydraulic conductivity, radial conductivity and/or axial conductance were determined experimentally. The Web of Science search engine was used for the review, and following search terms and keywords were included: *“root hydraulic conduct*”*AND *measur\** or *“root axial hydraulic conduct*”* AND *measur\** or *“root radial hydraulic conduct*”*AND *measur\**. The boolean operator AND was used to limit the search to studies in which root hydraulic properties were directly measured and not indirectly modelled from soil water content and/or plant transpiration or theoretically derived. All papers resulting from the search were revised in detail and only those which met the selection criteria were retained in the database.

In a second step, we checked the citations included in the selected papers to look for additional publications that may meet the selection criteria. Additionally, we looked at previous meta-analyses (Meunier *et al*., 2018; Bouda *et al*., 2018), reviews (Nobel & Cui, 1992; Huang & Nobel, 1994; Steudle, 2000a; North & Peterson, 2005; Maurel *et al*., 2010; Aroca *et al*., 2011; Gambetta *et al*., 2017; Kim *et al*., 2018) and the Xylem Functional Traits Database (Choat *et al*., 2012) to check for missing publications that should be included in our review. In total, we reviewed 241 papers, which comprises the vast majority of experimental studies on root hydraulic properties published between 1973-2023. A complete list of references included in the database is presented in Table S1.

### 2.2 Root hydraulic properties database

As part of the review process, we created an open access root hydraulic properties database, which aggregates all extracted data. Root hydraulic properties data were extracted manually and the software WebPlotDigitizer (Rohatgi, 2023) was used for digitalizing figures. The database contains detailed references to the original studies and provides easy, systematized access to the following data: root hydraulic properties (*K*_rs_, *k*_root_, *k*_r_ and/or *k*_x_), plant functional type (PFT, Table 2), growth form (a coarser classification than PFT, i.e. tree, shrub, succulent, graminoid and forb), tissue measured (whole root system, individual roots or root segments), root section (whole root or distal, mid-root or basal segments) measurement method, driving force for measurement, and experimental treatment(s) applied. When reported, plant age and morphological data were also included. The values stored in the database correspond to average values per study, species, factor (with factor being one or many among experimental treatment, tissue, root section, measurement method and driving force) and age. This means, for example, that a study reporting on *K*_rs_ of maize, based on two different measurement methods, with two treatments at three developmental stages generated a total of 1 × 2 × 2 × 3 = 12 data points. Therefore, the number of data points aggregated to the database from each study varied greatly. All digitized data is available for download in the database repository.

**Table 2:** Plant functional type (PFT) classification. . Selected PFTs and corresponding number of species, genera and studies for which root hydraulic properties were investigated. PFTs were defined based on commonly used classifications in land surface models (Poulter *et al*., 2015), and additional features such as growth form, differentiation between woody and herbaceous vegetation and agronomical importance.

| PFT | Description | Species examples | Nr. species | Nr. genera | Nr. studies |
| --- | --- | --- | --- | --- | --- |
| Crop herbaceous | Herbaceous crop species (legumes and non-legumes), excluding all C<sub>3</sub> and C<sub>4</sub> grasses | Tomato, soybean, lupin | 23 | 17 | 50 |
| Crop woody | Woody crop species | Cotton, grapevine | 2 | 2 | 11 |
| C<sub>3</sub> grass | Grass species with a C<sub>3</sub> photosynthetic pathway. Most species investigated corresponded to grasses used as crops | Barley, rice, wheat | 9 | 7 | 50 |
| C<sub>4</sub> grass | Grass species with a C<sub>4</sub> photosynthetic pathway. All species investigated corresponded to grasses used as crops | Maize, sorghum, pearl millet | 4 | 4 | 40 |
| Broadleaf tree | Deciduous and evergreen broadleaf tree species, including fruit trees | Quercus spp., Populus spp., Apple | 64 | 30 | 54 |
| Needle tree | Deciduous and evergreen needle tree species | Pinus spp., Picea spp., Abies spp. | 39 | 12 | 28 |
| Tropical tree | Broadleaf tree species from tropical ecosystems | Piper spp., Shorea spp. | 37 | 31 | 9 |
| Shrub | Deciduous and evergreen shrub species | Juniperus spp., Rhamnus spp. | 29 | 17 | 10 |
| Succulent | Succulent species from arid ecosystems | Agave spp., Opuntia spp. | 6 | 3 | 10 |
| Other | All species that could not be assigned to any of the defined PFTs | Arabidopsis thaliana., Dendrobium, Iris germanica | 3 | 3 | 6 |

Based on the digitalized data, we developed a web application (https://roothydraulic-properties.shinyapps.io/database/) that facilitates data selection, manipulation, visualization, and download. The main results presented in this study can be reproduced using the dynamic tools included there, and interested users are also encouraged to use these tools for their own research. The root hydraulic properties database, together with the web application, is conceived as a dynamic tool that will be updated continuously with newly reviewed studies. Readers are encouraged to share in the repository their new work or previously published work that may have been overlooked in our review process, by using the data sharing template available in the web application. The data included in the database is provided with free and unrestricted access for scientific (non-commercial) use (ODC-BY 1.0 license). Data users are requested to acknowledge the original data source and reference this review in resulting publications.

### 2.3 Data analysis and statistics

The data stored in the database was used for a comprehensive analysis on root hydraulic properties variability, excluding data that could not be classified into any PFT (defined as “Other”, see Table 2). The data was highly imbalanced, and there were large differences in the number of studies and species investigated for the different PFTs and root hydraulic properties. Accordingly, appropriate data analysis methods had to be selected. Although applying a strict meta-analysis (Hedges *et al*., 1999) could have been reasonable for this purpose, we discarded this approach because of two reasons: too few articles reported all the information needed for performing a meta-analysis (i.e., sample size and standard deviations for each experimental factor); and the experimental factors varied extremely among studies (Table S1), which hampered an evaluation of their individual effects and interactions. Instead, we followed an *ad-hoc* step-wise approach, and performed a series of independent analyses that quantified the variability in root hydraulic properties observed across studies and evaluated some of the (most important) factors causing it (see Table 3 for factor description). This analysis was performed for all individual root hydraulic properties except for *k*_r_, for which a very limited number of species and studies (*n*=12, in both cases) was available. Due to the large skewness in the original data, values were log transformed before data analysis, and then back transformed. Thus, the presented results correspond to geometric averages. Approximate standard deviations and standard errors were calculated using the Delta Method (Cramér, 1999).

**Table 3:** Factors affecting root hydraulic properties variability. Factors analyzed and their ranges (or factor levels) observed in the database.

| Factor | Description | Factor levels or range |
| --- | --- | --- |
| PFT | Plant functional types, according to the classification in Table 2 | Nine different PFTs |
| Age | Root system age.<br>Data principally corresponds to dicot crops and grasses. Root system age of trees and shrubs scarcely reported, mainly restricted to studies with seedlings | 3–150 days (herbaceous crops and grasses)<br>12–485 days (woody crops) |
| Driving force | Driving force used for measurement of root hydraulic properties | Hydrostatic or osmotic driving force |
| Genus | Taxonomic genus | 124 distinct genera |
| Growth form | A coarser classification than PFT | Tree, shrub, succulent, graminoid or dicot crops |
| Root section | Section of the root (segment) for which root hydraulic properties were determined. Several investigations measured whole roots instead of specific segments | Whole root or distal, mid-root or basal segments |
| Root type | Type of root investigated | Primary, tap, seminal, lateral, adventitious, whole root system |
| Species | Species investigated | 214 distinct species |
| Treatment | Simplified classification of the experimental treatments applied in the studies | Control, stress, other or no treatment |

In a first step, we calculated the range of variation (i.e., minimum, mean and maximum values) for each of the PFTs described in Table 2. For this, we first calculated the geometric means for the different studies and of each species investigated. These values were considered independent and suited for the analysis and were used for the calculation of the range of variation. The results corresponded to geometric means and range of variation for each PFT and root hydraulic property investigated (3.1).

Secondly, Random Forest (RF) models were run and the drop in accuracy of the model –a permutation feature importance metric (Altmann *et al*., 2010)– was calculated to rank the importance of several factors on the variability of root hydraulic properties. Next, linear mixed models were fitted to test for significant differences in root hydraulic properties among PFTs. PFT and two other highest ranked factors according to the RF model (excluding taxonomical features) were defined as the fixed effects, and study and experimental treatment were defined as the random effects. Given the extremely large dissimilarity in experimental designs among publications (see Table S1 for treatment list), we simplified the factor experimental treatment to four levels: control (defined as such in the publications), stress (any treatment that causes stress, e.g., drought, salt stress, nutrient limitation), other (any treatment that cannot be strictly defined as control or stress. e.g., different soil types, genotypes, season) and no treatment (studies where no treatments were applied). Type III ANOVA with the Satterthwaite’s method (Luke, 2017) was used for evaluating factor significance. The R-packages randomForest (Liaw & Wiener, 2002) and lme4 (Bates *et al*., 2015) were used for fitting the models.

Finally, we evaluated in more detail three factors that have been repeatedly reported to affect root hydraulic properties: driving force used for measurement, drought stress, and aquaporin (AQP) inhibition (see e.g., Aroca *et al*., 2011; Gambetta *et al*., 2017; Kim *et al*., 2018). For this, the natural log response ratio (ln(*r*) = ln(treatment) - ln(control)) (Hedges *et al*., 1999) was calculated for each individual study and species in which root hydraulic properties were measured under both treatment and control conditions. The results were reported as the mean percentage change ((*r* − 1)*100) (Ainsworth & Long, 2005) and response significance was tested with one-sample t-tests (on the log transformed data). Differences in the responses among PFTs were evaluated with one-way ANOVA tests. All data and statistical analyses were conducted in R v.4.3.1 (R Core Team, 2023).

### 2.4 Modelling the relationship between *K*_rs_ and root system age

The results of the RF and linear mixed models (see Section 3.2) indicated a significant and (probably) non-linear relationship between root system age and *K*_rs_ (and *K*_rs_area_). To investigate this relationship in more detail, we modeled the response of *K*_rs_ to the increase in root system age (and size) over time, using the functional-structural plant models CPlantBox (Schnepf *et al*., 2018) and MARSHAL (Meunier *et al*., 2019). Because data on root age was extremely scarce for trees and shrubs (see Table 3), this analysis was restricted to crop species (herbaceous crops and grasses).

CPlantBox was used to simulate the root system development of four different crops over a 120-day period: a C_3_ grass (wheat), a C_4_ grass (maize), a forb (cauliflower) and a legume (soybean). The species were selected based on plant-functional diversity and data availability. The XML-input parameters were obtained from the literature (Leitner *et al*., 2010; Vansteenkiste *et al*., 2014; Moraes *et al*., 2020; Morandage *et al*., 2021). CPlantBox outputs (i.e., the root architecture at each time step) were coupled to MARSHAL to simulate water flow from the soil-root interfaces to xylem vessels at the plant collar, using the analytical solution of water flow within infinitesimal subsegments (Meunier *et al*., 2017b), and to calculate the macroscopic parameter *K*_rs_ (Couvreur *et al*., 2012). Segment-scale *k*_r_ and *k*_x_ values were extracted from the database and from modelling (Doussan *et al*., 1998) and used to parametrize MARSHAL. *k*_r_ and *k*_x_ are age-dependent and vary among root types (Figure S1). To account for the uncertainty in their parameterization, a sensitivity analysis was performed by varying *k*_r_, *k*_x_ or the *k*_r_/*k*_x_ within the range of variation and the spatial heterogeneity observed in the literature (Figure S1). Modeled *K*_rs_ corresponds to the mean ± standard error of all simulations, for each individual crop. Modeling results were contrasted with data gathered from the review, specifically for crop species (dicot crops and C_3_ and C_4_ grasses) measured using a hydrostatic driving force.

## 3 Results and discussion

### 3.1 Range of variability of root hydraulic properties

In this work, we reviewed a total of 241 root hydraulic properties publications, comprising 215 species from 124 genera (complete list of references and species in Table S1). From this total, 165 studies focused on *K*_rs_, 60 on *k*_root_ (including *k*_r_) and 46 on *k*_x_ (some studies measured multiple hydraulic properties, simultaneously). We observed an extremely large range of variation (of orders of magnitude) in all root hydraulic properties, whereby this was especially pronounced for *K*_rs_ (Figure 2).

**Figure 2:**
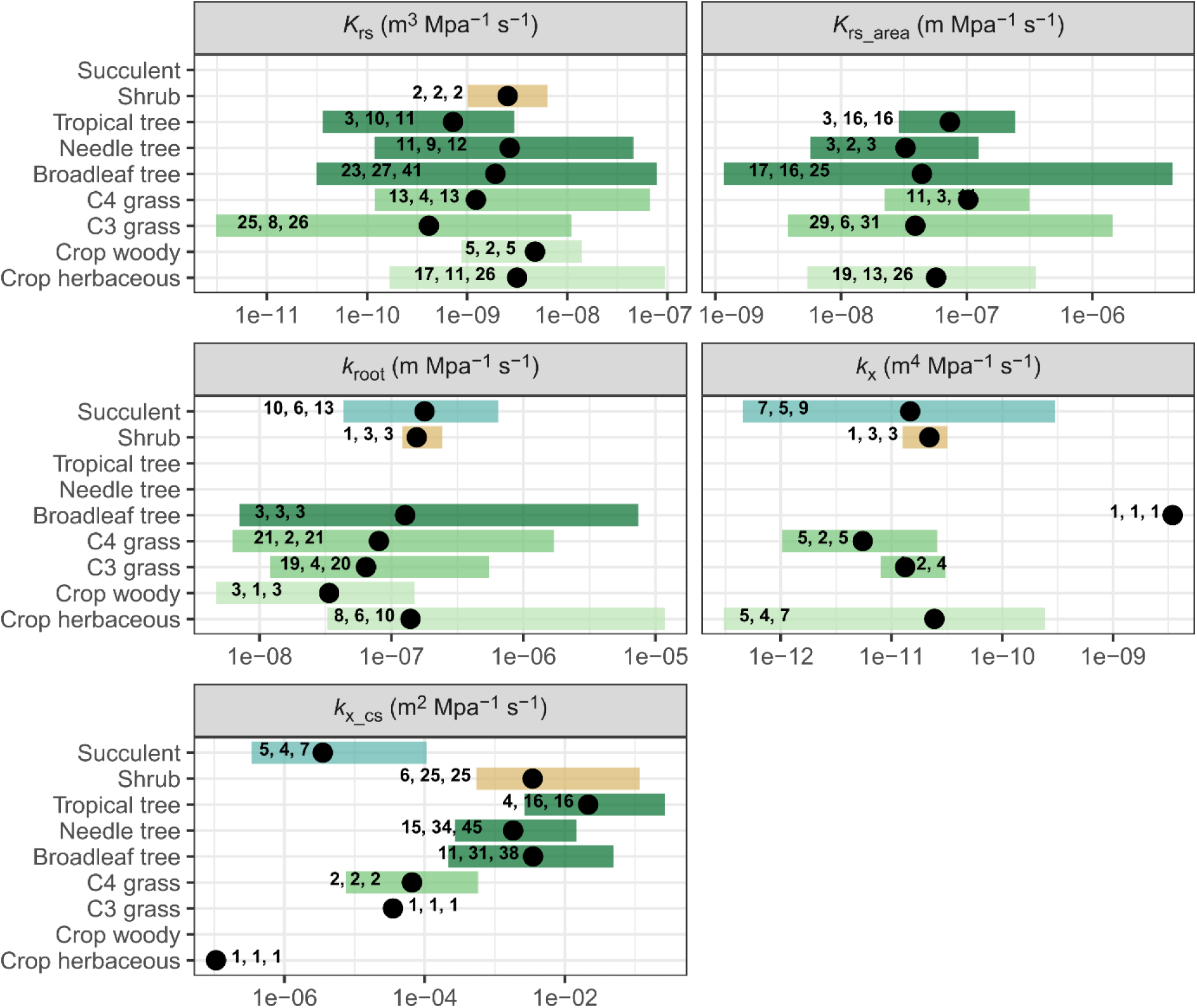
Range of variation in root hydraulic properties. Geometric means (filled circles) and range of variation (bars) of root hydraulic properties (see Table 1 for detailed definitions) for different plant functional types. The total number of studies, species, and individual data points for each PFT are indicated in bold (see 2.3 for details on the calculation).

Reported *K*_rs_ values varied extremely across studies, species, and plant functional types, ranging between 3.1×10^-12^ (measured in barley) to 9.4×10^-8^ m^3^ MPa^-1^ s^-1^ (measured in common bean). A very large range of variation was also observed within PFTs, with *K*_rs_ showing a range of variation of ≈ 2–3 orders of magnitude in all PFTs, except for shrubs (for which only two studies were available). This was considerably larger than the differences in the geometric means among PFTs, which varied between 4.1×10^-10^ (C_3_ grasses) and 4.8×10^-9^ m^3^ MPa^-1^ s^-1^ (woody crops). Due to the very large intra-PFT variability, possible systematic differences among PFTs could have been obscured (but see 3.2.1).

*K*_rs_ is often reported in the literature on the basis of a measure of root size, to facilitate the comparison among plants of different age, with root surface area (*K*_rs_area_) being the normalization most widely used (see Table 1 for other common normalizations). Our results indicated that the range of variation of *K*_rs_area_ was indeed factors of magnitude smaller than that of *K*_rs_, but it was still extremely large (1.2×10^-9^ – 4.3×10^-6^ m MPa^-1^ s^-1^) (Figure 2). A very large range of variation was also observed within each PFT (≈ 1–3 orders of magnitude), indicating large intrinsic differences among species and/or experimental design of the studies. Surprisingly, even, both the lowest and the highest *K*_rs_area_ values found in the literature corresponded to broadleaf tree species (*Q. petraea* and *P. tremula × tremuloides*). On the contrary, the geometric mean of *K*_rs_area_ varied comparatively slightly among PFTs (3.3×10^-8^ – 1.0×10^-7^ m MPa^-1^ s^-1^).

Published root hydraulic properties data of individual roots and/or root segments (total, radial, and axial) also showed very large variability. The total conductance *k*_root_ (which is often reported as a proxy of *k*_r_ in the literature) varied extremely across studies (range = 4.7×10^-9^ – 1.2×10^-5^ m MPa^-1^ s^-1^, Figure 2), but also within individual PFTs (ranges ≈ 1–3 orders of magnitude). This large variation was observed despite the few species that have been investigated (2–6 species for the different PFTs). Additionally, the geometric means of *k*_root_ showed small variation among PFTs (3.4×10^-8^ –1.8×10^-7^m MPa^-1^ s^-1^), and this range was almost identical to that of *K*_rs_area_.

Axial conductance also showed a very large variability, both for published data reported as *k*_x_ (range = 3.1×10^-13^ – 3.5×10^-9^ m^4^ MPa^-1^ s^-1^) and on a cross sectional area basis (*k*_x_cs_, range = 1.1×10^-7^ – 2.7×10^-1^ m^2^ MPa^-1^ s^-1^). However, we found very few studies on *k*_x_ (20 publications), and they were unevenly distributed across PFTs. While succulent species were the most frequently reported (7 studies, 5 species), only one tree species was available and showed by far the largest *k*_x_ (1-3 order of magnitudes larger than any other value). Excluding that species, *k*_x_ ranged between 3.1×10^-13^– 3.0×10^-10^ m^4^ MPa^-1^ s^-1^, with C_4_ grasses showing the lowest (5.5×10^-12^ m^4^ MPa^-1^ s^-1^) and dicot crops the highest (2.4×10^-11^ m^4^ MPa^-1^ s^-1^) geometric means among PFTs. At the same time, *k*_x_cs_ has been widely reported for woody vegetation (26 publications, 105 species) and showed a range of variation between 2.2×10^-4^ – 2.7×10^-1^ m^2^ MPa^-1^ s^-1^, with tropical trees showing the highest values. These values were systematically higher than *k*_x_cs_ of the very few non-woody species for which data was available (8 species, range = 1.1×10^-7^ – 1.1×10^-4^ m^2^ MPa^-1^ s^-1^) and confirm the results from previous metanalyses (Bouda *et al*., 2018). However, our review also highlights the difficulty of comparing axial conductance of woody and non-woody vegetation, with the former almost entirely being reported as *k*_x_cs_ and the latter as *k*_x_.

### 3.2 Understanding root hydraulic properties variability

The results in Section 3.1 showed an extremely large range of variation in root hydraulic properties across published studies. Here, we further investigated to which degree the observed variability could be explained by the response of root hydraulic properties to the following factors: systematic differences among PFTs, driving force used for measurement (hydrostatic or osmotic), effect of environmental stresses, and root system age.

#### 3.2.1 Main factors affecting root hydraulic properties and differences among PFTs

One central question we addressed in this study was whether the observed variability in root hydraulic properties could be attributed to systematic differences among PFTs. For this, we first used Random Forest (RF) regressions to compare the importance of PFT with other variables that have been reported to affect root hydraulic properties. This included factors such as root system age, the driving force used for measurement (hydrostatic or osmotic), root section and root type, experimental treatment, or variation within species. According to the “drop in accuracy” metric (Table 4, more details in 2.3), root system age had the highest importance to explain the variability in *K*_rs_, which agrees with the general positive relationship between *K*_rs_ and root system size observed in the literature (Tyree, 2003). This is the case, as with increasing age the root system grows, adding conductances (new root segments) in parallel in a hydraulic network, which increases the total conductance of that network. Interestingly, root system age also showed the highest importance for *K*_rs_area_, suggesting complex interactions between root system growth and *K*_rs_ development (see 3.2.4 for further discussion). The importance of PFT for *K*_rs_ was 27.4% smaller (and 26.9% smaller for *K*_rs_area_) than that of root system age and was similar to the importance of driving force or species and only clearly larger than that of experimental treatment (Table 4). These results indicate that the large variability of *K*_rs_ observed in the literature cannot be explained by systematic differences among PFTs, alone, but rather by the added effect of multiple factors.

**Table 4:** Statistics of Random Forest and linear mixed models. . Importance of several factors (as described in Table 3) for root hydraulic properties variability, according to the drop in accuracy metric (Random Forest); *p*-value of the same factors, using Type III ANOVA tests (linear-mixed models); and total variance explained by the fitted Random Forest models. Data in bold indicate the 3 highest ranked factors (Random Forest models) and effect significance (p < 0.05, ANOVA tests).

| Factor | Drop in mean square error |  |  |  | *p*-value (Satterthwaite) |  |  |  |
| --- | --- | --- | --- | --- | --- | --- | --- | --- |
|  | *K<sub>rs</sub>* | *K<sub>rs_area</sub>* | *k<sub>root</sub>* | *k<sub>x_cs</sub>* | *K<sub>rs</sub>* | *K<sub>rs_area</sub>* | *k<sub>root</sub>* | *k<sub>x_cs</sub>* |
| PFT | **2.07** | 0.98 | **0.91** | **7.39** | 0.20 | 0.84 | 0.92 | **<0.001** |
| Age | **2.85** | **1.34** | - | - | **<0.001** | 0.38 |  |  |
| Driving force | 1.63 | **1.1** | **0.95** | - | **<0.001** | **<0.001** | **<0.001** |  |
| Genus | 1.84 | 1.05 | **0.89** | 1.81 | - | - | - | - |
| Growth form | 1.14 | 0.41 | 0.68 | **6.72** | - | - | - | - |
| Root section | - | - | 0.64 | **2.47** |  |  | **<0.01** | **<0.001** |
| Root type | - | - | 0.75 | 1.79 | - | - | - | - |
| Species | **1.99** | **1.12** | 0.76 | 1.8 | - | - | - | - |
| Treatment | 0.67 | 0.33 | 0.31 | 0.41 | - | - | - | - |
|  | - | - | - | - | - | - | - | - |
| **Total variance explained (%)** | 76.9 | 65.9 | 64.3 | 83.6 | - | - | - | - |

We also analyzed the importance of PFT for *k*_root_ (Table 4) and observed that it was lower than the importance of driving force (−4.2 %) and slightly higher to that of species, root type (seminal, adventitious, lateral) or root section (distal, mid-root, basal or entire root). This suggests that the observed variability of *k*_root_ is caused by the added effect of multiple factors and their interactions, rather than by systematic differences among PFTs. However, care must be taken in the interpretation of these results, due to the rather small number of species investigated (26) and the extremely low number of studies (5) in which species belonging to different PFTs were investigated simultaneously. On the contrary, the importance of PFT for *k*_x_cs_ variability was much larger (at least more than twice) than that of any other factor, except for growth form, confirming the clear, systematic difference between woody and non-woody species depicted in Figure 2 and the observations of Bouda *et al*. (2018). These results are probably associated with large increases in axial conductance (2-3 orders of magnitude) following secondary growth in woody roots (Vercambre *et al*., 2002) and with large differences in xylem cross sections between woody and non-woody vegetation.

To confirm the results of the RF models and further investigate systematic differences in root hydraulic properties among PFTs, individual linear mixed models for *K*_rs_, *K*_rs_area_, *k*_root_ and *k*_x_cs_ were run, with PFT and additional non-taxonomical features (i.e. root system age, driving force, root section or root type, detailed factor and model description in Section 2.2–2.3) as fixed effects, and study and treatment as random effects.

We found no significant effect of PFT on *K*_rs_ (*p* = 0.20), *K*_rs_area_ (*p* = 0.84) and *k*_root_ (*p* = 0.92), but *k*_x_cs_ varied highly significantly (*p* < 0.001) among PFTs (Table 4), which agrees with the results of the RF analysis and its conclusions. On the contrary, a highly significant effect of driving force (*p* < 0.001) on *K*_rs_, *K*_rs_area_ and *k*_root_ was found, indicating systematic difference in root hydraulic properties measured using a hydrostatic driving force, against those using an osmotic driving force (see 3.2.2 for a detailed analysis). Additionally, root system age showed a highly significant positive effect on *K*_rs_ (*p* < 0.01), probably associated with an increase of *K*_rs_ with increasing root system size. Conversely, root system age had no effect on *K*_rs_area_ (*p* = 0.38), contradicting the high importance that root age had for *K*_rs_area_ prediction, according to the RF model. Interestingly, though, the linear mixed model showed a negative (albeit non-significant) relationship between *K*_rs_area_ and root age and this negative relationship became significant (*p* < 0.05) when a negative exponential function was fitted to the data, instead of a linear relationship. This implies a decrease in *K*_rs_ per unit root surface over time, a phenomenon that could be associated with the decrease in segment-scale radial conductivity with age, but also with axial transport limitation with increasing root length (Meunier *et al*., 2017b; Bouda *et al*., 2018, see also discussion in Section 3.2.4). Clearly, the relationship between root age and *K*_rs_ (and *K*_rs_area_) observed in our review is complex and was therefore explored in more detail in Section 3.2.4.

The linear mixed models also showed a highly significant (*p* < 0.001) effect of root section –a factor describing whether root hydraulic properties were measured on basal, mid-root or distal root segments or on entire roots– on *k*_root_ and *k*_x_cs_, suggesting the presence of spatial gradients in roots across species and PFTs. Spatial variation alongside roots in *k*_r_ and *k*_x_ (and consequently in *k*_root_) has been reported for the grass species maize (Frensch & Steudle, 1989; Doussan *et al*., 1998; Meunier *et al*., 2018) and barley (Knipfer & Fricke, 2011) and for *A. deserti* (Huang & Nobel, 1992), with radial conductivity decreasing from root tip to root base, while the opposite was the case for axial conductance (see also Figure S1). Variation can be caused by changes in root anatomy and function (e.g., formation of apoplastic barries, increase in xylem diameter and density, differences in aquaporin expression) with increasing age. However, similar gradients were not evident (particularly in the case of *k*_r_ and *k*_root_) in onion (Melchior & Steudle, 1993) or lupin (Doussan *et al*., 2006; Meunier *et al*., 2018), questioning the idea that they are ubiquitous across species and PFTs. Our review cannot answer this, because most of the studies reported data for one root section only, hampering systematic comparison among sections. For instance, the two largest *k*_root_ values in our review (1.2×10^-5^ in *V. faba* and 7.4×10^-6^ in *P. trichocarpa x deltoides*) corresponded to measurements in distal segments, but unfortunately no other root section was investigated in those studies. Nevertheless, the statistical results underscore the significance of spatial gradients as a factor of variability in root hydraulic properties and stress the need for further investigations on this topic, focusing on the differences (or lack thereof) among species from different PFTs.

In general, the statistical analyses did not reveal systematic differences in root hydraulic properties among PFTs, apart from the highly significant effect of PFT on axial conductance, a feature that has been reported previously. Rather, the results imply that the variation in multiple factors such as age, driving force, or root section analyzed (and probably their interactions) determined the extremely large variability observed here. This would also explain why root hydraulic properties varied so much within PFTs (Figure 2) or even within species. Accordingly, a detailed analysis on the influence of several factors on root hydraulic properties variability (with the main focus on *K*_rs_) was also performed in this review, and the results are presented in the following sections (3.2.2 – 3.2.4).

To our knowledge, this is the first systematic review on the topic of root hydraulic properties and their variability across PFTs, leaving little room for the comparison of our results with previous investigations. However, we cannot discard the possibility that systematic differences among PFTs –which we did not find– were obscured by the dissimilarity in experimental design among the publications. Actually, less than 10% of the reviewed studies included species corresponding to more than one PFT, and the hydraulic properties investigated there were unevenly distributed: while *K*_rs_ and *k*_root_ studies mostly focused on dicot and monocot crop species (Gallardo *et al*., 1996; Bramley *et al*., 2007; e.g. Hess *et al*., 2015), broadleaf and needle trees were predominant in *k*_x_ (or rather *k*_x_cs_) studies (e.g. Maherali *et al*., 2006; Domec *et al*., 2010). In fact, we only found one study in which root hydraulic properties of trees and herbaceous vegetation were measured simultaneously (Rieger & Litvin, 1999). Thus, more studies comparing root hydraulic properties across species and PFTs are needed to confirm (or reject) the results in this review.

#### 3.2.2 The driving force matters

According to the results from the previous section, the driving force used for measurement was a key factor for explaining the very large variability observed in this review. Here, we quantified in more detail the differences in root hydraulic properties (specifically *K*_rs_ and *k*_root_; *k*_x_ data is not relevant for this analysis) estimated under osmotic gradients (hereafter osmotic root hydraulic properties), compared to those estimated under hydrostatic gradients (hereafter hydrostatic root hydraulic properties), based on the log response ratio of pairwise comparisons (methodological details in 2.3).

A total of 39 data pairs, corresponding to 29 studies and 16 species were investigated, whereby only four species (maize, barley, rice, and wheat) accounted for >60% of all values (see Table S2 for all studies and species included). On average, osmotic root hydraulic properties were 78.1% smaller than hydrostatic ones, and this effect was highly significant (*p* < 0.001). More interestingly, the observed response varied significantly among PFTs (*p* < 0.001), showing average decreases ranging from 42.6% (C_3_ grasses) to 94.9% (broadleaf trees). In that, C_3_ grasses showed a much lower decrease compared to the remaining PFTs, which varied very slightly among each other (range = 94.9 – 85.4%; woody crops were not included in this comparison, because only one value was available). For all PFTs, the reported decrease in osmotic root hydraulic properties (Figure 3) was significantly different from zero (*p* < 0.05).

**Figure 3:**
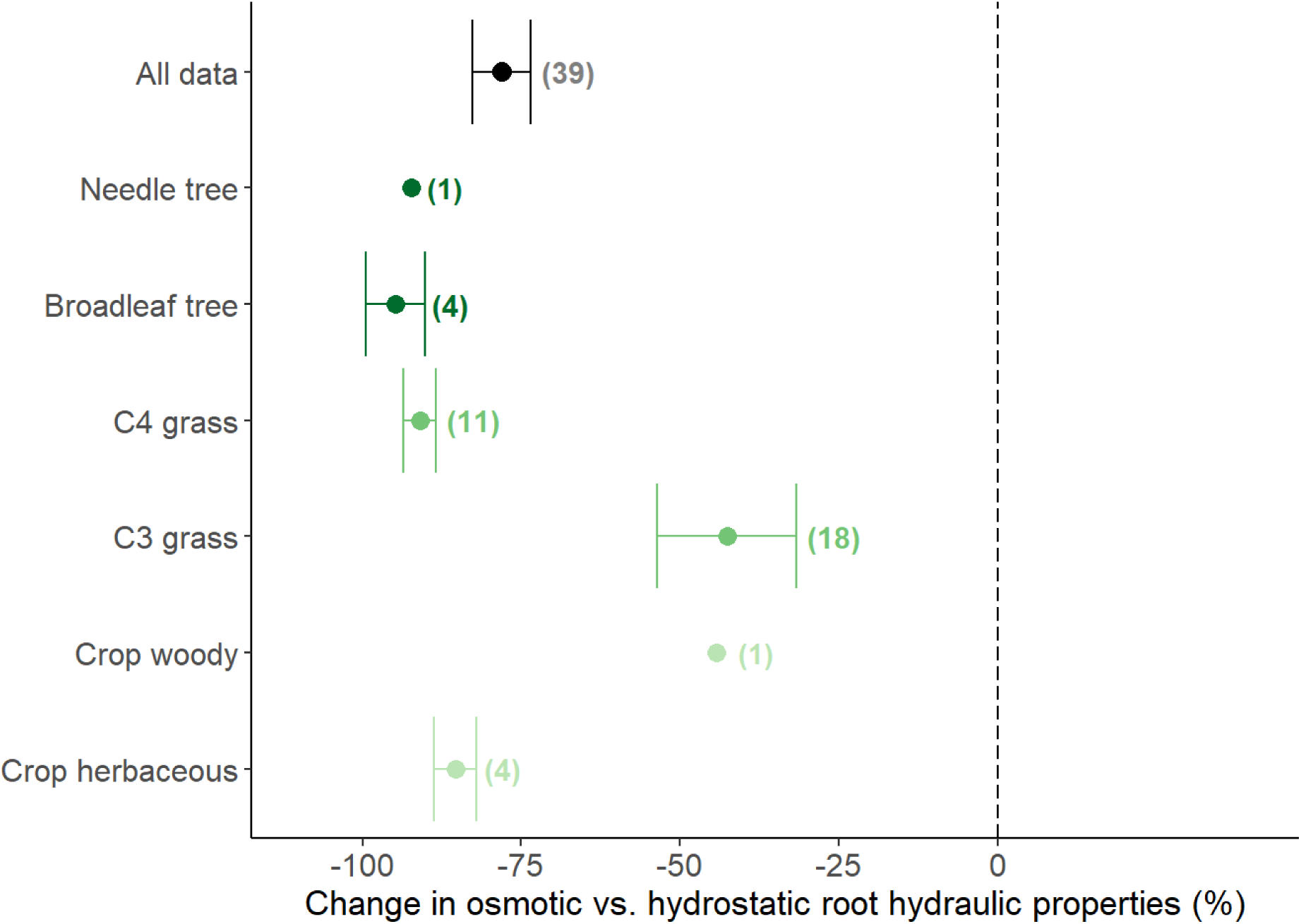
Difference between osmotic vs. hydrostatic root hydraulic properties. Data points and error bars represent the mean ± the standard error for each PFT (sample size *n* reported on the side). The mean value for all samples is represented with a black circle. Individual values were calculated based on the log response ratio.

Clearly, the driving force affects the measurements of root hydraulic properties. Across all studies, the largest difference was observed in *K*_rs_ of oak trees and reached almost two orders of magnitude (Steudle & Meshcheryakov, 1996). On average, a decrease of ≈78% of osmotic compared with hydrostatic root hydraulic properties was observed, and in four PFTs (broadleaf and needle trees, C4 grasses and dicot crops) a decrease of ≈90% (i.e., 1 order of magnitude) was reached. Considering that the total range of variation within PFTs was ≈1–3 orders of magnitude (Figure 2), the driving force can be described as one of the most important factors for explaining the variability in root hydraulic properties reported in this review.

That osmotic root hydraulic properties are systemically lower than hydrostatic ones has been reported before (Steudle, 2000a; Kim *et al*., 2018). In line with the principles of the composite transport model (Steudle, 2000a), the comparison between osmotic and hydrostatic root hydraulic properties has been widely used to differentiate the cell-to-cell path (obtained from osmotic measurements) from the overall path for water flow (i.e. cell-to-cell + apoplastic paths, obtained from hydrostatic measurements) and how the contribution of the former might change under conditions of environmental stress (see e.g. Garthwaite *et al*., 2006; Barrios-Masias *et al*., 2015; Kreszies *et al*., 2020). According to this approach, our results would imply that the cell-to-cell path had a (much) smaller contribution than the aploplastic path to the total water flow across PFTs, with the cell-to-cell contribution to total water flow being the lowest in broadleaf trees (4.9%) and the highest in C3 grasses (36.5%). However, the accuracy of this approach has been questioned (Chaumont & Tyerman, 2014), as multiscale studies do not support this common assumption and rather indicate that the differences between osmotic and hydrostatic root hydraulic properties may stem from an erroneous estimation of the osmotic driving pressure and therefore of hydraulic properties (Bramley *et al*., 2007; Couvreur *et al*., 2018). Cell-scale simulations of the advection-diffusion of osmolytes suggest that their accumulation at apoplastic barriers (e.g. Casprian strip) may alone generate a 5-fold overestimation of the effective water potential gradient across the endodermis (Knipfer & Fricke, 2011, Steudle, 2008; Couvreur *et al*., 2018), while apoplastic, symplastic and transmembrane modes of water transport would vary radially regardless of whether the water potential difference between root surface and xylem is due to pressure or osmolytes. Nevertheless, the data clearly showed a differentiation between C_3_ grasses and the remaining PFTs, and also very large discrepancies within the C_3_ grasses: while osmotic and hydrostatic root hydraulic properties were almost equal in barley (≈6% higher osmotic root hydraulic properties, in average), osmotic root hydraulic properties were much smaller than hydrostatic ones in wheat and rice (≈55% and ≈63% in average, respectively). To which degree these differences indicate functional heterogeneity in water transport patterns among species lies beyond the scope of this review, but the data presented here could be used to identify species or PFTs of interest for future studies.

#### 3.2.3 Responses to drought and AQP inhibition

Environmental stress has been widely reported as a factor affecting root hydraulic properties (Steudle, 2000b; Maurel *et al*., 2010; Aroca *et al*., 2011; Gambetta *et al*., 2017). Interestingly, though, our analysis showed that experimental treatment had the lowest importance of all variables in explaining the range of variation in *K*_rs_, *K*_rs_area_, *k*_root_ and *k*_x_cs_ observed in the literature (Table 4). Two aspects could explain these results: (1) the variation across studies and PFTs was so large, that it obscured the effects of experimental treatments observed in individual studies; and (2) experimental treatments differed extremely among studies (Table S1), hindering a systematic analysis of the effect of environmental stress on root hydraulic properties variability. Thus, for the purpose of this review, the response of root hydraulic properties to stress was narrowed to two factors: drought stress and aquaporin (AQP) inhibition. For this, 28 studies on the effect of drought stress and 19 studies on the effect of AQP inhibition on *K*_rs_ (or its normalized values) were analyzed.

There was a significant decrease in *K*_rs_ under both drought stress and AQP inhibition (*p*<0.001 in both cases). On average, *K*_rs_ decreased 61% under drought conditions and the decrease under AQP inhibition was very similar (59%). However, the *K*_rs_ response to drought showed more variation across PFTs, studies or species than that to AQP inhibition. The average *K*_rs_ decrease under drought varied among PFTs in a range between 80.8% (in dicot crops) and 38.3% (in C_3_ grasses), and this variation was marginally significant (*p* = 0.07) (Figure 4). Meanwhile, *K*_rs_ decreased under AQP inhibition in a smaller range between 50.9% (in tropical trees) to 77.4% in (C4 grasses) (*p* = 0.16). Also, across all studies and species (*n*=30), the *K*_rs_ response to drought varied greatly, between ≈98% decrease (i.e., a decline of almost two orders of magnitude) and ≈35% increase. On the contrary, *K*_rs_ responded negatively to AQP inhibition, without exception (*n*=25), with the decrease ranging between ≈22%–86%.

**Figure 4:**
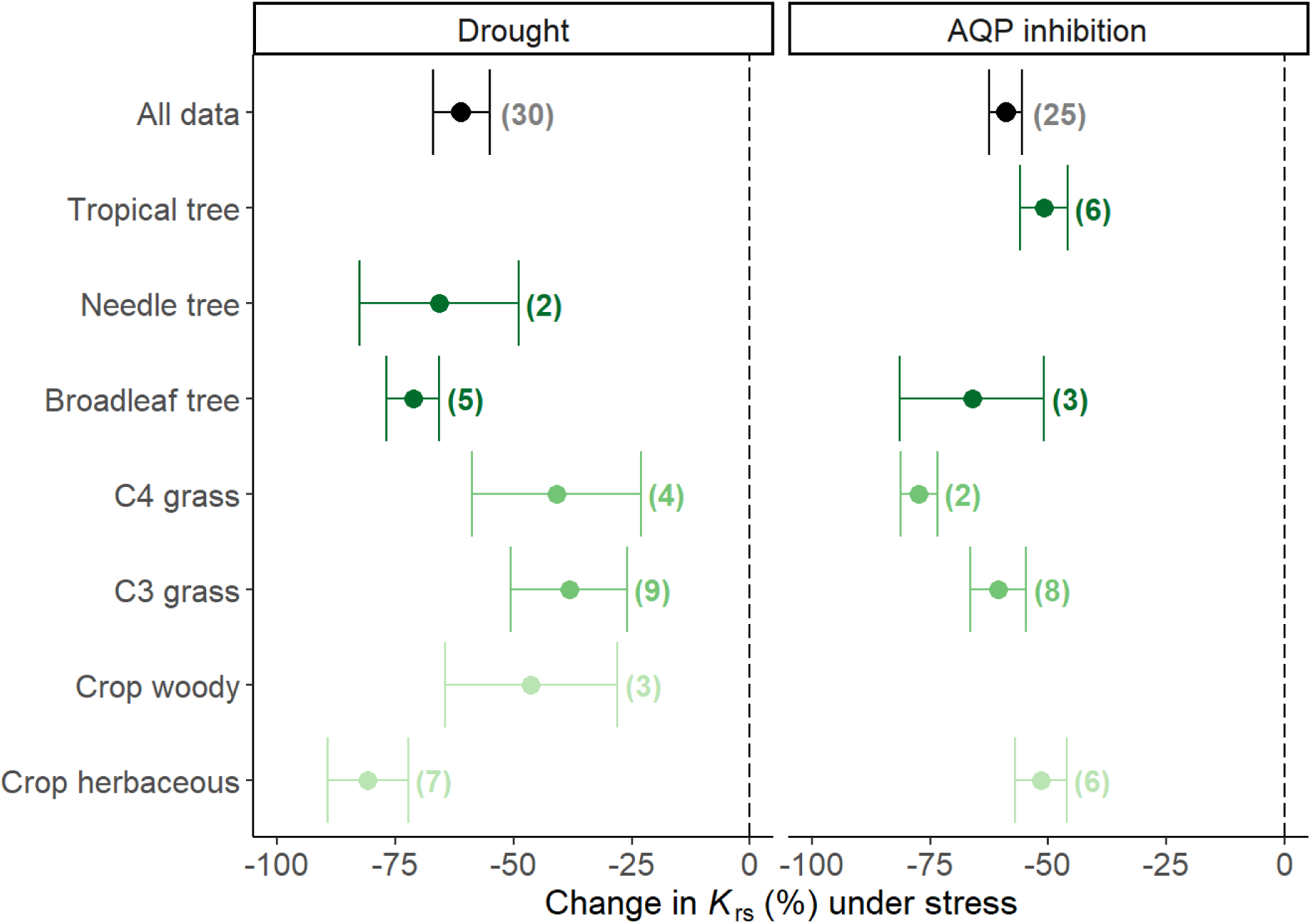
Response of *K*_rs_ to stress treatments. Changes in *K*_rs_ under drought stress (left panel) and aquaporin inhibition (right panel). Data points and error bars represent the mean ± the standard error for each PFT (sample size *n* reported on the side). The mean value for all samples is represented with a black circle. Individual values were calculated based on the log response ratio.

The average decline in *K*_rs_ under drought agrees with the conclusions of previous reviews (Aroca *et al*., 2011). This response corresponds to a water saving strategy under condition of limited water availability, which can be induced by short-term responses (e.g., changes in the aquaporin gating), but also on long-term drought-driven anatomical changes (e.g., formation of apoplastic barriers, aerenchyma, changes in xylem vessel size) or changes in root size (Aroca *et al*., 2011; Vadez, 2014; Bauget *et al*., 2023). Furthermore, our review revealed differences among PFTs (albeit non-significant, probably due to a small sample size), with grasses (both C_3_ and C_4_) showing a weaker response to drought than trees or dicot crops. In fact, the only three studies in which an increase in *K*_rs_ under drought was reported, were conducted with rice (Lian *et al*., 2004; Ding *et al*., 2015) and maize (Zhang *et al*., 1995). Also, the *K*_rs_ decrease of maize (C_4_ grass, ≈44%) under drought was considerably weaker than that of tomato (dicot crop, ≈63%), in the only study where grass and non-grass species were directly compared (Bárzana *et al*., 2012), supporting the overall trends reported here. However, the shown differences among PFT might be conditioned by the low number of species investigated within each PFT. For example, in the case of C_3_ grasses seven out of 9 studies were conducted with rice, and a similar behavior was observed for C_4_ grasses (all 4 studies with maize) or dicot crops (4 out of 7 studies with tomato). But, regardless of these limitations, our results contribute to a better understanding of the expected root hydraulic properties variability under drought conditions across species and PFTs.

On the other hand, a negative response of *K*_rs_ to AQP inhibition was observed across all PFTs and species investigated. This effect is driven by a decrease in the cell-to-cell radial water flow (Aroca *et al*., 2011; Chaumont & Tyerman, 2014), such that the large range in *K*_rs_ responses to AQP inhibition (≈22%–86% decrease across studies) could be associated with differences in aquaporin activity of root cells among the investigated species and PFTs. However, we did not observe systematic differences among PFTs in our analysis. In a previous review on aquaporins and root water uptake, Gambetta *et al*. (2017) also identified a very large range in the response of root hydraulic properties to AQP inhibition, and mainly attributed this to variability in the experimental approach across studies. As such, further examinations of the responses exhibited by distinct tissues, species, and/or plant functional types (PFTs) are essential to enhance our understanding of water flow dynamics under stress conditions, and how this might impact the overall variability of root hydraulic properties.

#### 3.2.4 Non-linear K_rs_ increase with increasing root system age in crops and grasses

Root system age is a key factor for explaining the large variability in *K*_rs_ observed in this review (see 3.2.1). Here, we investigated this relationship in more detail, for hydrostatic *K*_rs_ of dicot crops and grass species (selection criteria described in 2.4). Across studies and species, there was a significant increase in *K*_rs_ with increasing age of the root system (*p* < 0.01), with the relationship exhibiting a non-linear pattern (Figure 5). *K*_rs_ increased abruptly during the first 20–30 days of root development, and then slowly flattened out, with a total range of variation between ≈6×10^-11^ – 2×10^-8^ m^3^ MPa^-1^ s^-1^. The steep increase in *K*_rs_ during the first days of development is probably caused by the growth of the root system adding new conductances (new roots) to the root hydraulic network, thus increasing the total conductance of the network. However, the asymptotic behaviour after days 30-40 suggests a partial decoupling between root size and *K*_rs_ at later stages of development. Unfortunately, root size data (e.g., root surface area or total root length) was not reported ubiquitously across studies, impeding the analysis of the interactions between *K*_rs_, root age and root size. Interestingly, though, an analogous asymptotic relationship between root length and *K*_rs_ has been previously reported in a modelling study (Meunier *et al*., 2017a).

**Figure 5:**
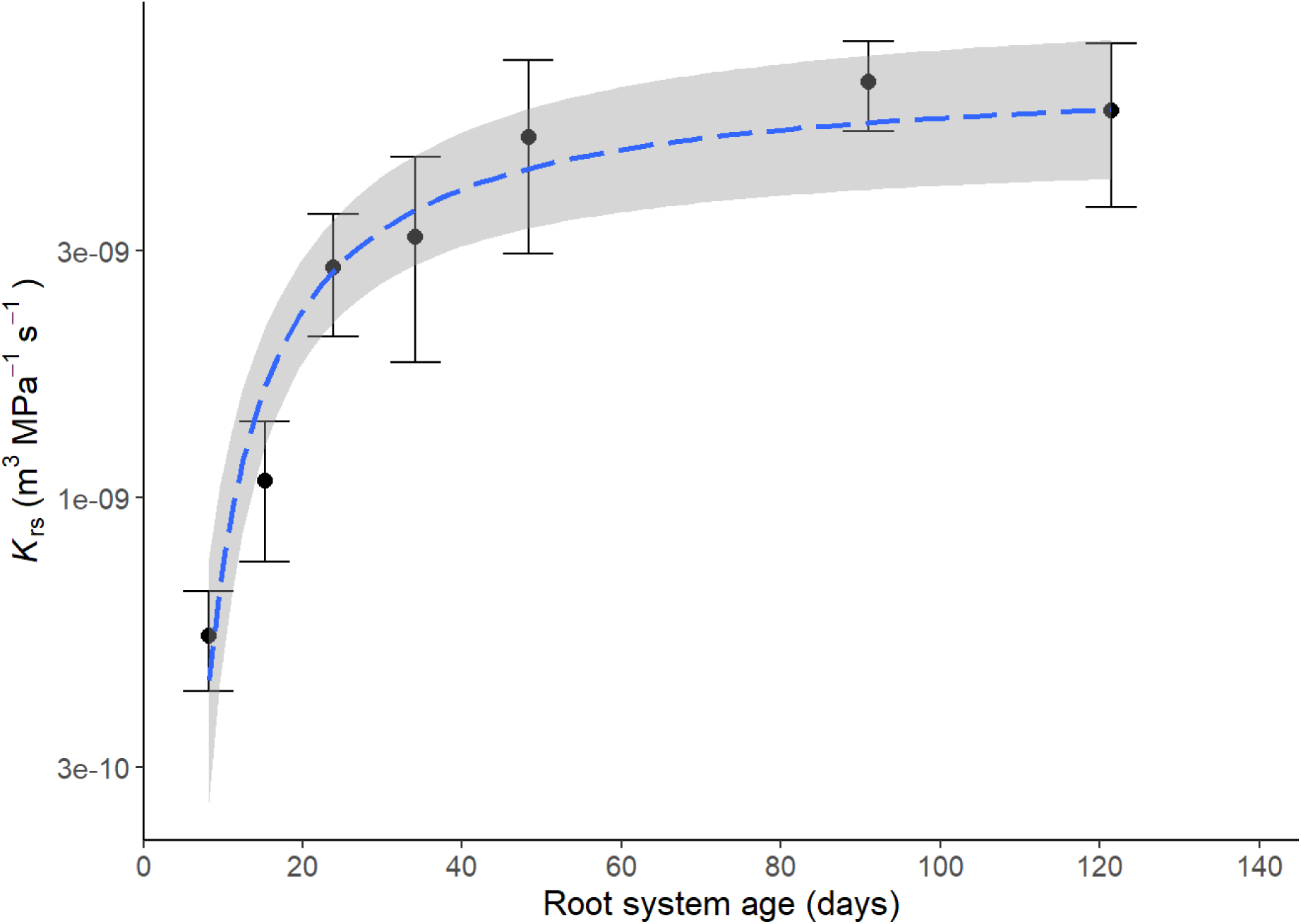
Relationship between root system age and *K*_rs_. Data points and error bars represent *K*_rs_ (mean ± standard error) of crop species grouped according to age (0–10, 10–20, 20–30, 30–40, 40–60, 60–100, >100 days). The dashed blue line and the shaded area represent a fitted exponential model (± standard error).

To explore the *K*_rs_ development with age in more detail, we modeled this relationship for four selected crop species, using CPlantBox coupled with MARSHAL (see 2.4 for details on data selection and model parametrization). Despite large differences in root size and root architecture (Figure S2), all species exhibited a very similar non-linear pattern, i.e., a pronounced increase in *K*_rs_ with age during the first 20 days, followed by rather constant values from day 20 onwards (Figure 6). This behaviour was not related to cessation in root growth, as total root length showed a continuous increase during the 120 days of simulation (Figure S2). But, with increasing root age the proportion of “old” root segments (> 10-day old segments) also increased (Figure 6). This could have impacted the development of *K*_rs_, as the radial (*k*_r_) and axial (*k*_x_) hydraulic properties of root segments –which, together with the root architecture, determine *K*_rs_– are age dependent (Doussan *et al*., 1998). Specifically, *k*_r_ strongly decreases with age (Figure S1), and the radial pathway is commonly considered to be the more limiting one for water transport (Frensch & Steudle, 1989; Lynch *et al*., 2014). Thus, the counteracting effect of an increase in less conductive tissues (i.e., older root segments) proportionally to total root growth would explain the constancy in *K*_rs_ at later stages of development. Additionally, it has been shown that even under constant *k*_r_ and *k*_x_, *K*_rs_ can display an asymptotic behavior for roots due to axial flow limitations with increasing root length (Meunier *et al*., 2017a). Furthermore, the modeled *K*_rs_ response to age strongly resembled the one observed in the empirical data. In fact, average *K*_rs_ values at different ages obtained from the review lay within (or very near) the range of variation of the models (Figure 7), indicating that the modelling results were representative of common patterns across studies and species. Whether the mechanisms observed in the models also explain the patterns evidenced in the review remains to be investigated.

**Figure 6:**
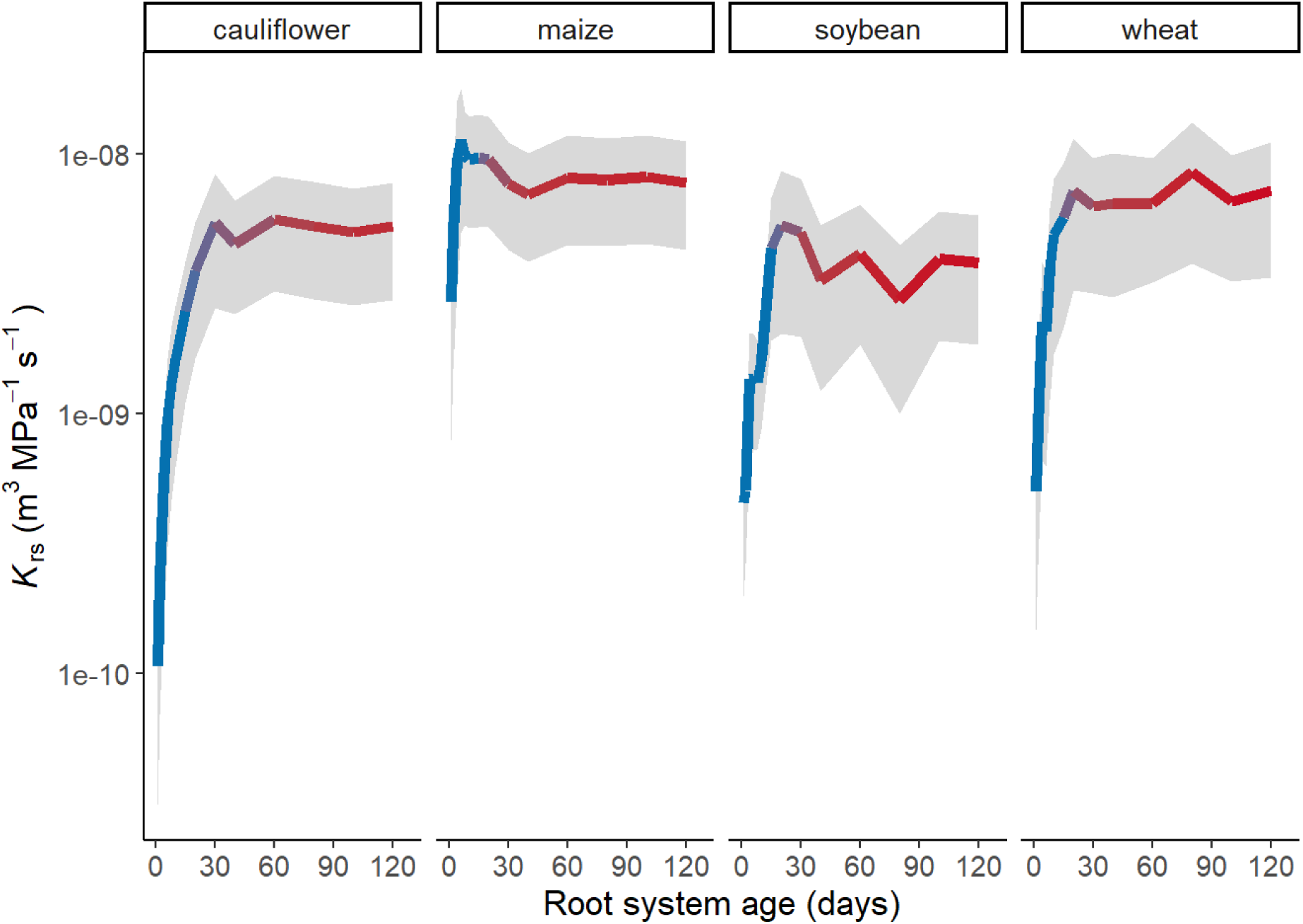
Modelled *K*_rs_ development with age. Colored lines and shaded areas represent *K*_rs_ (mean ± standard error) of simulations using CPlantBox coupled with MARSHAL, for four different crops. The color scale indicates the proportion of old (>10 days) root segments in the total root system.

**Figure 7:**
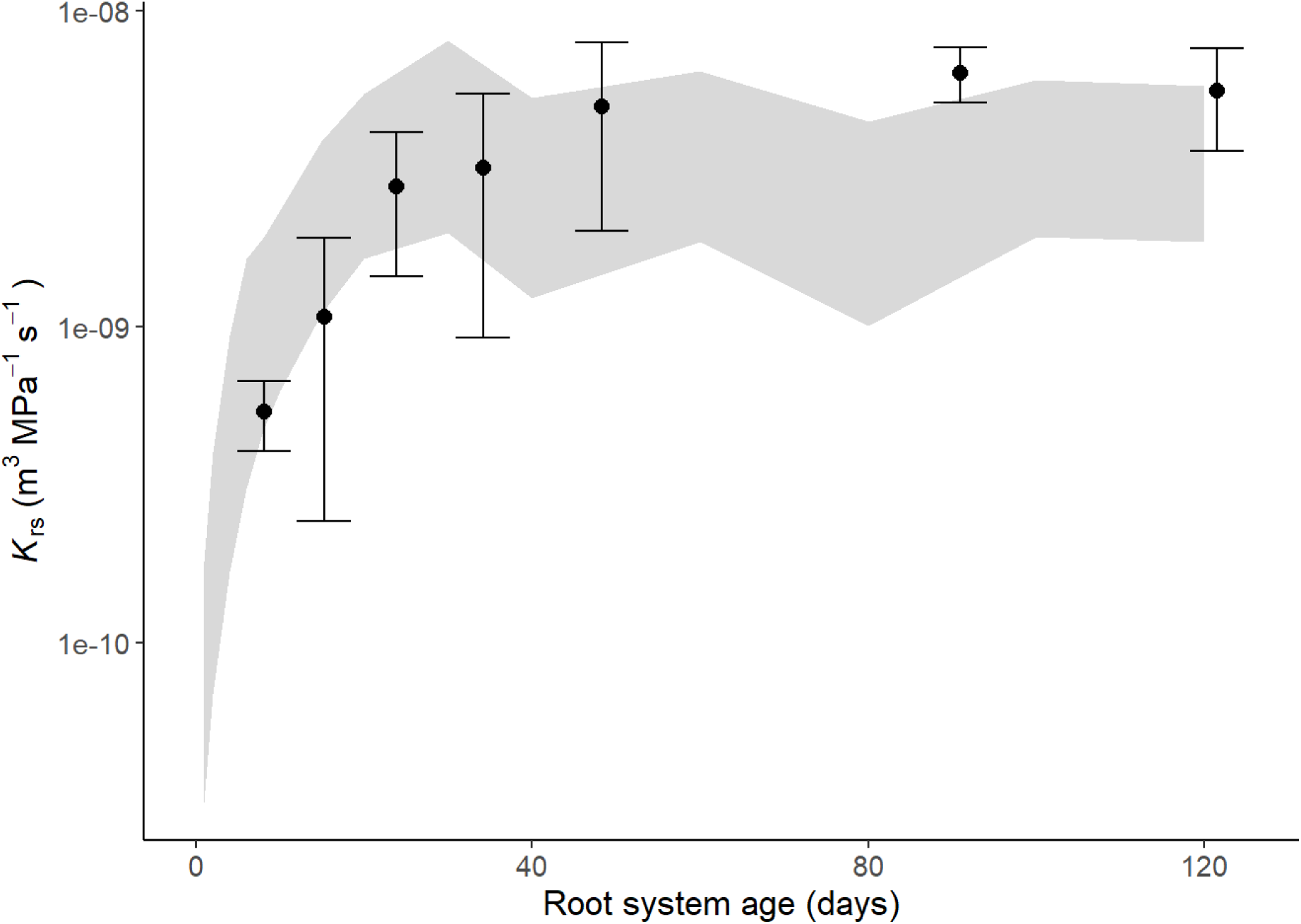
Modelled and observed *K*_rs_ development with age. Data points and error bars represent *K*_rs_ (mean ± standard error) of crop species from the review and the shadowed area represents the total range of variation in *K*_rs_ according to simulations (CPlantBox coupled with MARSHAL).

The non-linear relationship between *K*_rs_ and root system age presented here has been reported previously. For instance, a similar pattern was observed in a modelling study with 10,000 virtual maize root systems (Meunier *et al*., 2019). However, our work is the first –at least to our knowledge– to demonstrate a common pattern across studies and species in both experimental data and modelling and to quantify the associated range of variation in *K*_rs_ over time. Also, the combination of literature data and modelling gave insights about the (possible) causes for the emerging patterns. These results are therefore of relevance and can be a valuable input for the description of root water uptake processes at plant, field or regional scales (Couvreur *et al*., 2014; Sulis *et al*., 2019; Nguyen *et al*., 2020; Vanderborght *et al*., 2021; Nguyen *et al*., 2022; Jorda *et al*., 2022).

## 4 Conclusions and outlook

Here, we presented an extensive review on root hydraulic properties, their variability and some of the factors affecting them. A very large range of variation (orders of magnitude) in *K*_rs_, *k*_root_, *k*_r_ and *k*_x_ reported in the literature was identified, but this was not caused by systematic differences among plant functional types (with the only exception of significant differences between axial conductance of woody vs. non-woody species), but rather by the (combined) effect of factors such as root system age, driving force used for measurement, root tissue measured, environmental stress or intra-specific variation. As a result, a closer examination was undertaken to explore the influence of some of these factors on root hydraulic properties. This yielded new insights on root hydraulic properties variability, some of which could not be analyzed here in detail, due to the inherent limitations of a broad review, but should be targeted specifically in future studies. The following topics are of special interest: (1) the difference between osmotic and hydrostatic root hydraulic properties was much lower in C_3_ grasses (particularly in barley) than in other PFTs; how is this reflected in the water transport patterns of these species?; (2) a large range of variation was observed in the response of root hydraulic properties to drought, with some indications of differences among PFTs, but clear conclusions were hindered by the extremely low number of studies comparing multiple species and PFTs. Hence, do species corresponding to different PFTs (e.g. dicot crops vs. grasses) respond differently to drought under the same environmental conditions?; and (3) a common non-linear relationship between root system age and *K*_rs_ was identified for several crop species, according to both literature data and modelling. Is such a pattern also present in species from other PFTs (e.g., shrubs or young trees) and how is it reflected in the seasonality of perennial species?

In summary, the present study represents an overview of root hydraulic properties variability across plant functional types, species and experimental conditions and their associated responses. The new insights obtained here, together with the accompanying data (stored in a database and easily accessible through the web application, https://roothydraulic-properties.shinyapps.io/database/) and additional tools like modelling –as we applied in this study– should be a valuable input for future studies on the role of root hydraulics and root water uptake processes under changing environmental conditions.

## Author contributions

JCBC: Conceptualization, Software, Formal Analysis, Investigation, Data Curation, Visualization, Writing - original draft, Writing - review and editing

JV: Formal Analysis, Funding Acquisition, Writing - review and editing

VC: Formal Analysis, Writing - review and editing

DB: Writing - review and editing

TG: Funding Acquisition, Writing - review and editing

THN: Writing - review and editing

GL: Formal Analysis, Funding Acquisition, Writing - review and editing

## Acknowledgements

This research was supported by the Deutsche Forschungsgemeinschaft (DFG, German Research Foundation), in the DETECT - Collaborative Research Center (SFB 1502/1-2022 - Projektnummer: 450058266). VC is a Research Associate of the Belgian Fonds National pour la Recherche Scientifique (FNRS), co-funded by the European Union (ERC grant 101043083). THN is part of the “COINS project”, funded by the Federal Ministry of Education and Research (BMBF).

