## Supplemental Figures 1-2 for "Root hydraulic properties: an exploration of their variability across scales"

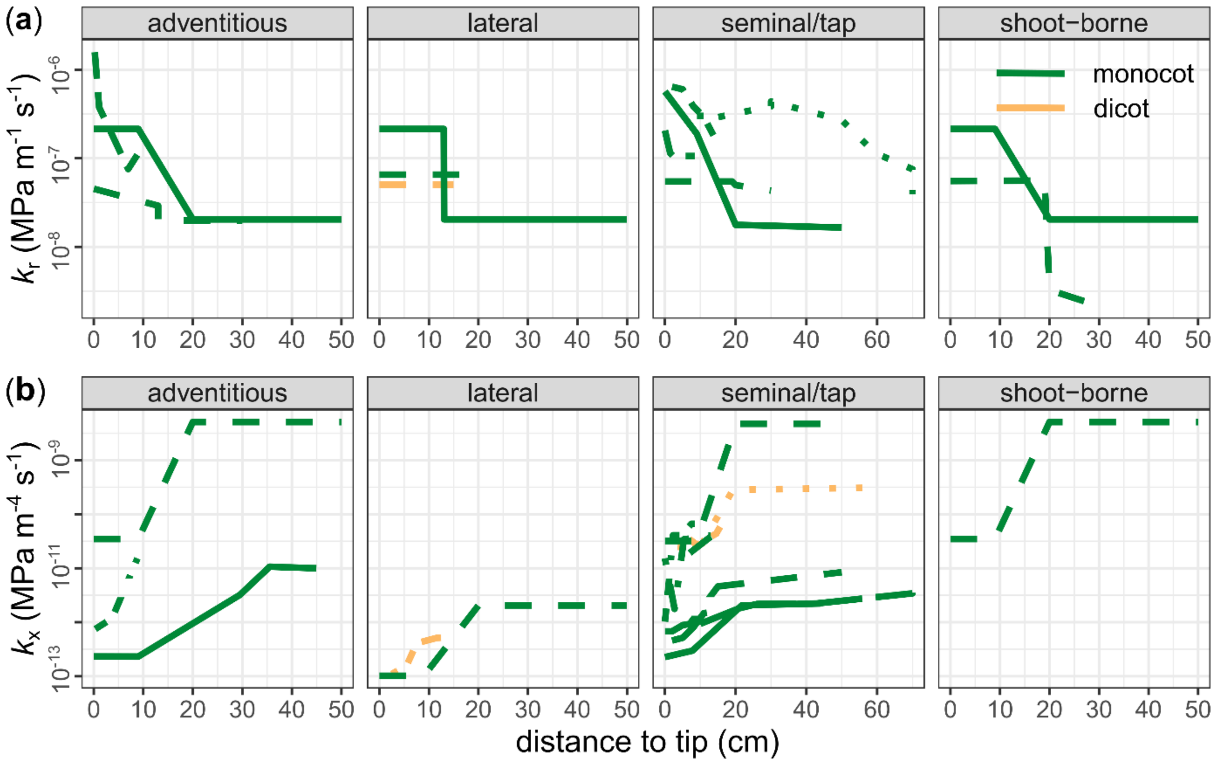


**Figure S1**: **Spatial gradients in root hydraulic properties**. Spatial variation in radial conductivity ($k_r$) (**a**) and axial conductance ($k_x$) (**b**) alongside roots (from tip to base), for different root types in monocot (green) and dicot (yellow) herbaceous species. The data corresponds to published studies on segment-scale root hydraulic properties (Ahmed *et al*. 2018; Bramley *et al*. 2007; Bramley *et al*. 2009; Doussan *et al*. 1998; Doussan *et al*. 2006; Frensch & Steudle, 1989; Huang & Steudle, 1992; Knipfer & Fricke, 2011; Meunier *et al*. 2018; Nobel *et al*. 1993; Ranathunge *et al*. 2017). The different line types indicate different references.

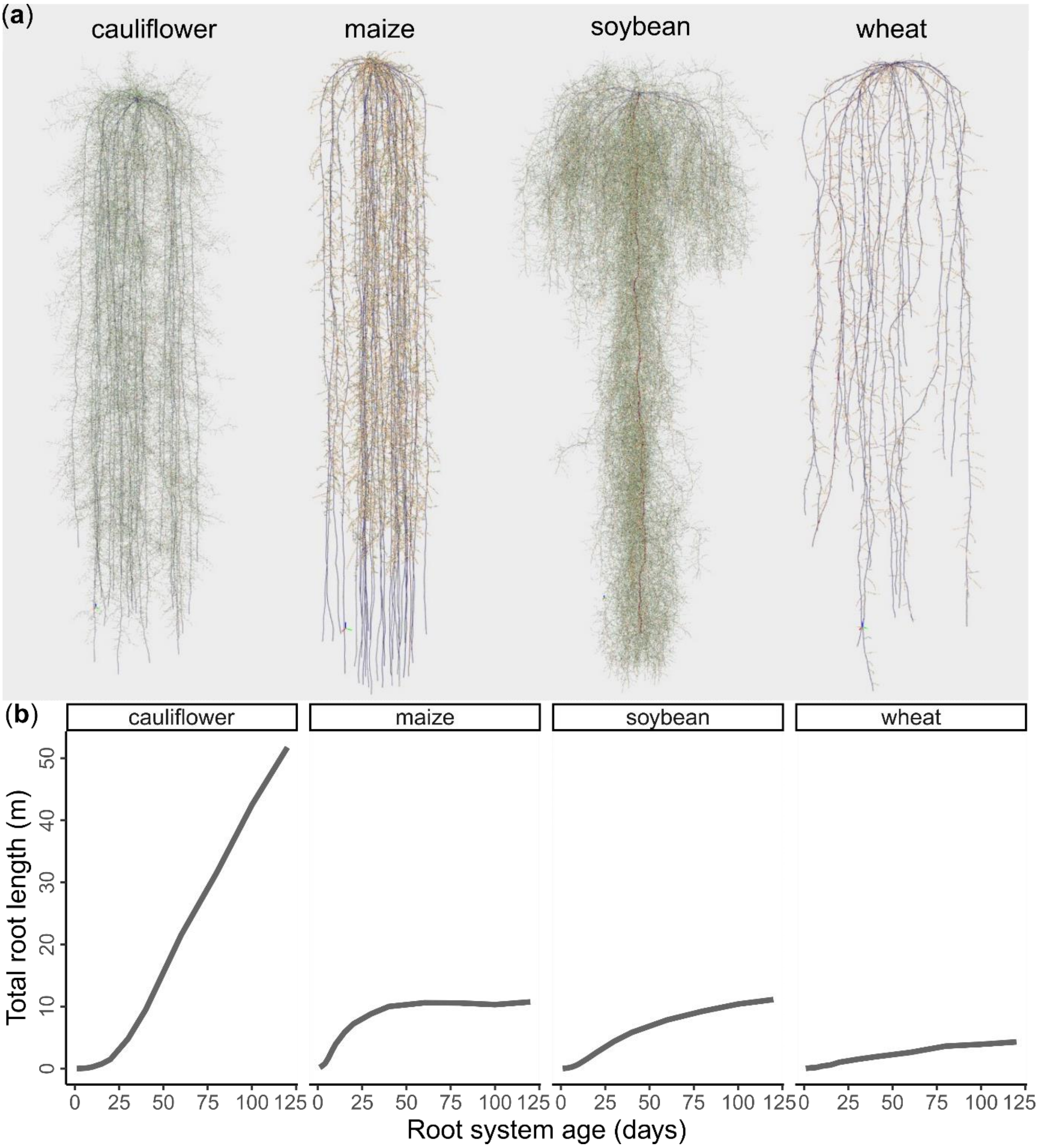


**Figure S2: Root architecture and root size of different crops**. Root architecture at day 120 (**a**) and total root length development during a 120-day period (**b**) for four different crop species: cauliflower (dicot herbaceous crop), maize ($C_4$ grass), soybean (legume crop) and wheat ($C_3$ grass). Root architecture development was simulated with CPlantBox (Schepf *et al*. 2018), based on available XML-files from the literature (Leitner *et al*. 2010: Moraes *et al*. 2021; Morandage *et al*. 2021; Vansteenkiste *et al*. 2014).
